# Two new plasmid post-segregational killing mechanisms for the implementation of synthetic gene networks in *E. coli*

**DOI:** 10.1101/350744

**Authors:** Alex J H Fedorec, Tanel Ozdemir, Anjali Doshi, Luca Rosa, Oscar Velazquez, Tal Danino, Chris P Barnes

## Abstract

Plasmids are the workhorse of both industrial biotechnology and synthetic biology, but ensuring they remain in bacterial cells is a challenge. Antibiotic selection, commonly used in the laboratory, cannot be used to stabilise plasmids in most real-world applications, and inserting dynamical gene networks into the genome is difficult. Plasmids have evolved several mechanisms for stability, one of which, post-segregational killing (PSK), ensures that plasmid-free cells do not grow or survive. Here we demonstrate the plasmid-stabilising capabilities of the axe/txe two component system and the microcin-V system in the probiotic bacteria *Escherichia coli* Nissle 1917 and show they can outperform the hok/sok system commonly used in biotechnological applications. Using plasmid stability assays, automated flow cytometry analysis, mathematical models and Bayesian statistics we quantified plasmid stability *in vitro*. Further, we used an *in vivo* mouse cancer model to demonstrate plasmid stability in a real-world therapeutic setting. These new PSK systems, plus the developed Bayesian methodology, will have wide applicability in clinical and industrial biotechnology.

## Introduction

The genes comprising a synthetic circuit can be maintained in a host bacterium in two ways: on the chromosome of the organism or on extra-chromosomal material such as plasmids. Plasmids are a fundamental biological tool and have been widely used in molecular and cellular biology research, leading to a number of well developed methods for their manipulation (Ellis et al. 2011, Casini et al. 2015). This ease of manipulation enables a level of modularity which is one of the key engineering goals of synthetic biology (Andrianantoandro et al. 2006, Martinez-Garcia et al. 2014). Plasmid copy number can be an important parameter in the functioning of circuits, and it is non-trivial to convert a dynamical gene circuit from a multi-copy plasmid implementation to a single copy implementation and maintain identical function (Lee et al. 2016).

However, one of the fundamental problems with using plasmids is their segregational instability. When bacteria divide there is the possibility that all plasmid copies remain in one half of the cell, which leads to the production of a plasmid-free daughter cell, as shown in Figure 1A. As most of the synthetic circuits that are borne on plasmids produce a burden to their hosts, the plasmid-free population outgrows the plasmid-bearing population and the engineered strain is quickly diluted from the environment (Summers 1991). Maintaining the presence of the engineered circuit within the bacterial population is fundamentally important in the design of a predictable synthetic biological system.

**Figure 1:**
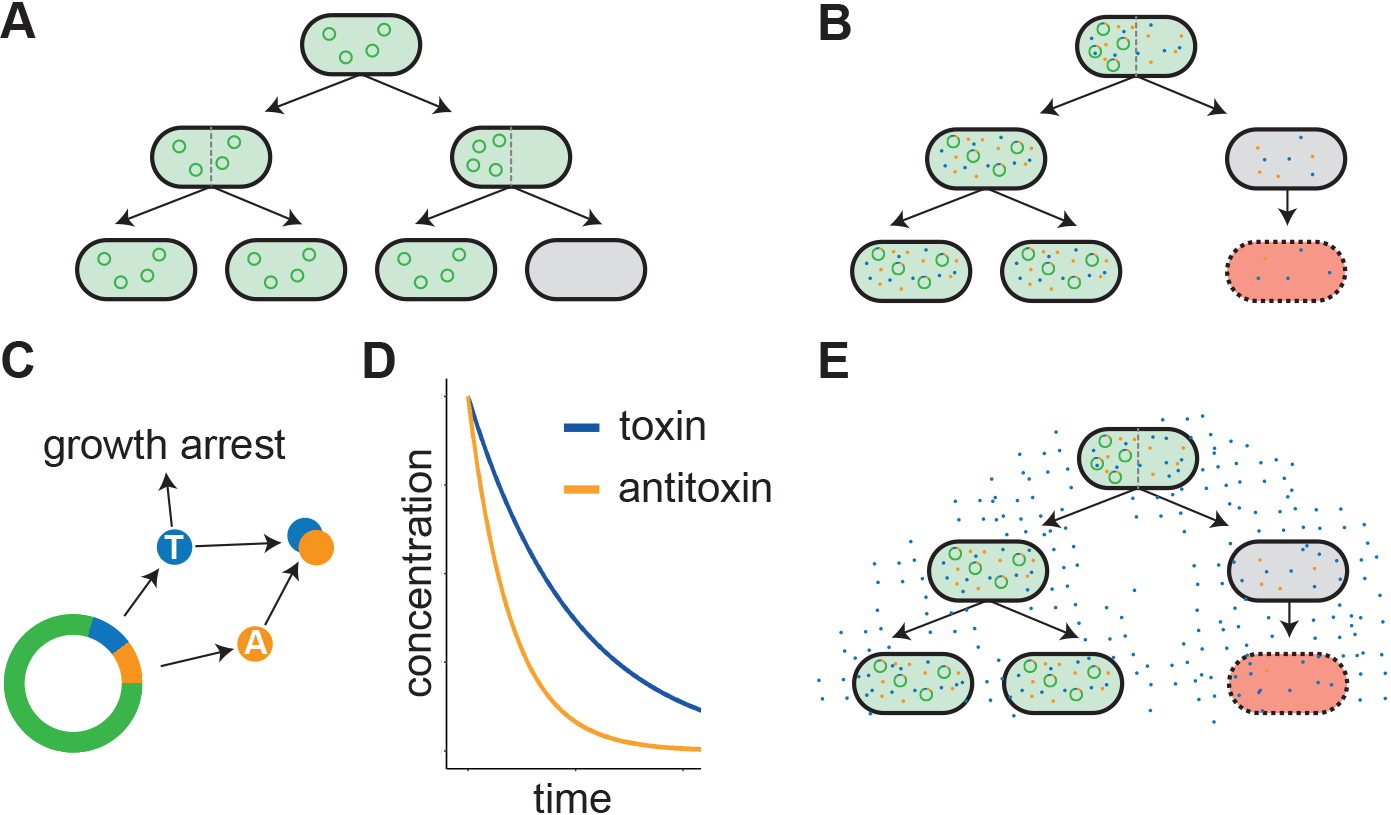
Plasmid-stability and post-segregational killing. A) plasmid free cells are produced due to uneven distribution of plasmids at cell division. B) Post-segregational killing mechanisms carried on plasmids cause the death of cells that lose the plasmid when dividing. C, D) The toxin-antitoxin PSK systems are plasmid-borne mechanisms that rely on a long-lived toxin degrading more slowly in a newly plasmid-free cell than the antitoxin. E) Bacteriocins can improve plasmid stability by a population level policing through the secretion of a toxin in to the environment.

Antibiotic selection of bacteria containing plasmids carrying the corresponding antibiotic resistance genes is commonly used in a research environment. However, antibiotics are not used in industrial fermentation due to the financial impact of removal and deactivation (Kroll et al. 2010). Antibiotics are also unsuitable for clinical applications for a number of reasons including horizontal gene transfer of resistance genes (Stecher et al. 2012, Wright et al. 2008) and the disruption of the native microbiota (Theriot et al. 2014). In light of these limitations, efforts have already been made to re-use a variety of existing microbial machinery to ensure plasmid persistence in more complex environments (Kroll et al. 2010, Wright et al. 2013). Successful alternatives have been demonstrated with the use of toxin-antitoxin (TA) systems (Loh & Proft 2013, Danino et al. 2015), active-partitioning mechanisms (Danino et al. 2015, Liu et al. 2005) and auxotrophy (Velur Selvamani, Friehs & Flaschel 2014, Velur Selvamani, Telaar, Friehs & Flaschel 2014).

TA systems are a type of post-segregational killing mechanism that work through the production of a long lasting toxin and its shorter lived antitoxin (Cataudella et al. 2012, Diago-Navarro et al. 2013), both encoded on the plasmid, as in Figure 1B,C. While the plasmid is present in the cell, antitoxin is being produced to neutralise the toxin. If, however, a plasmid-free daughter cell arises, the antitoxin quickly degrades and is unable to be replaced due to lack of the necessary genes. The negative effects of the toxin are no longer prevented, leading to the killing or growth prevention of the cell (Figure 1D). The successful post-segregational killing of a plasmid-free cell is reliant on enough toxin becoming active in the cell before it goes on to divide again, further diluting the toxin. Just as with the distribution of plasmids giving a probability of producing plasmid-free cells, there is a probability that the distribution of toxin is such that the TA system fails to kill the newly plasmid-free cell. As such, TA systems are fallible and once plasmid-free cells escape, they have no mechanism for preventing their growth and the dilution of the plasmid-bearing population. Hok/sok is a type I TA system originating from the *parB* locus of the *E. coli* plasmid R1 and has been shown to be effective in its native *E. coli* as well as the gram-negative *Pseudomonas putida* (Gerdes 1988). Axe/txe is a proteic, type II TA system originating from the *axe*-*txe* locus of the gram-positive *Enterococcus faecium* plasmid pRUM (Grady & Hayes 2003). It has been demonstrated that axe/txe could be used to stabilise a luminescent reporter in the gram-positive *Enterococcus faecalis in vivo* without antibiotic selection for five days (Rosa et al. 2013). The axe/txe system was also found to be present in 75% of vancomycin-resistant enterococci (VRE) isolates, a common hospital pathogen and growing global concern (Moritz & Hergenrother 2007).

An alternative to TA systems are bacteriocins, which are bacterially secreted proteins that have a bactericidal affect on either a narrow or broad spectrum of other bacteria lacking immunity (Riley 1998). By secreting these antimicrobial peptides, a plasmid-bearing population is able to police the environment, preventing the growth of plasmid-free cells (depicted in Figure 1E). Successful attempts have already been made to use bacteriocins, such as the Lcn972 system, to stabilise plasmids in *L. lactis* (Campelo et al. 2014) and the colicins A and E2 in *E. coli* (Inglis et al. 2013). In comparison to the TA system, bacteriocins can maintain plasmids even with mutational instability. Whereas a single mutation in the toxin on a TA plasmid would remove the selective pressure in that cell, the selection with bacteriocins is maintained via the surrounding cells. Regardless of whether they are able to produce the bacteriocin, each cell still needs to encode the immunity gene on the plasmid to avoid death (Inglis et al. 2013).

Here we use the microcin-V system which is encoded on conjugative plasmids in *E. coli*. It consists of a low molecular weight toxic bacteriocin which kills through pore formation, along with the genes for an ABC-transporter and immunity protein (Azpiroz & Lavia 2007). It does not undergo any post-translational modification and, as such, does not require any extra enzyme encoding genes. The structure of the bacteriocin gene itself is also modular, which allows for its hybridisation with other bacteriocins, enabling the targeting of other bacterial strains (Acuña et al. 2012, 2015). Further, it has been shown that EcN already carries two bacteriocin systems in its chromosome; microcins H47 and M (Grozdanov et al. 2004). More recently, it was also shown that native bacteriocins found in EcN were vital in mediating competition in the inflamed gut in a inter-and intraspecies manner (Sassone-Corsi et al. 2016).

In this work we develop a mathematical model and Bayesian inference procedure to quantify the efficacy of two PSK systems, axe/txe and microcin-V, in stabilising a fluorescent reporter plasmid in EcN and compare them to the more commonly used and studied hok/sok system. The model enables determination of post-segregational killing efficacy and also the extra metabolic burden due to the PSK system. We also investigate the ability of the PSK systems in stabilising a luminescent reporter plasmid without antibiotic selection *in vitro* and in a mouse tumour xenograft model *in vivo*. Collectively, we show that the two new systems show a much greater potential for plasmid stability than hok/sok in EcN.

## Results

### A mathematical model for post-segregational killing mechanisms

Early mathematical models of plasmid stability consisted of terms for a plasmid-bearing population and plasmid-free population, growing exponentially at different rates with a constant probability of plasmid loss from the plasmid-bearing population (Boe et al. 1987). These models were then used to determine plasmid loss rates from experimental data (Boe 1996, Boe & Rasmussen 1996) and extended to describe bacterial populations in a chemostat; introducing a rate of dilution as well as growth rates dependent on a substrate (Ganusov & Brilkov 2002). A more detailed model of plasmid loss was devised that takes into account the age distribution of bacteria within a population, although the predictions produced were virtually indistinguishable from simpler models (Lau et al. 2013). However, this model did highlight the difficulties of trying to infer the plasmid loss rate from measured population data as, due to the exceedingly small plasmid loss rates of most systems, the growth rate differences dominate the dynamics and lead to over-estimation of plasmid loss (Lau et al. 2013).

Here, we extend the early plasmid loss model (Boe 1996) to include post-segregational killing by a toxin-antitoxin system. In this model there are two populations; plasmid-bearing (*X*^+^) and plasmid-free (*X*^−^)

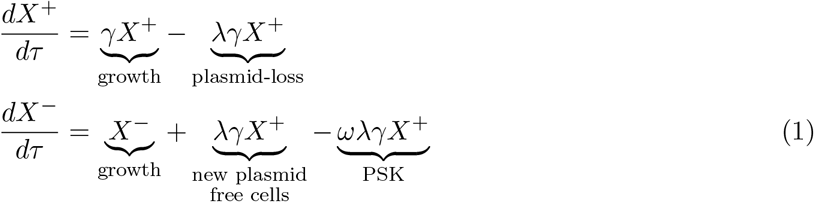

where λ is the probability of producing a plasmid-free daughter cell when a plasmid-bearing cell divides, γ is the ratio of plasmid-free doubling time to plasmid-bearing doubling time, and ω is the probability of successful post-segregational killing. The time-step τ is equal to one plasmid-free generation.

Bacteriocins do not produce post-segregational killing in the traditional sense. Instead there is a constant killing pressure on the plasmid-free population rather than specific killing of newly plasmid-free cells. As such, the equation for the change in plasmid-free population differs to take into account the different point at which the bacteriocin causes the death of plasmid-free cells.

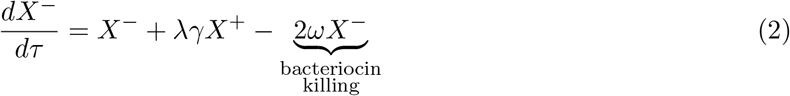

Supplementary Figure 1 shows the effects of varying each of the model parameters on the dynamics of plasmid loss. The plasmid loss parameter, λ, affects the gradient of the plasmid loss curves early on, when there are very few plasmid-free cells. As soon as a plasmid-free population is established, the doubling time ratio, γ, has a large effect on the gradient. For the TA model in Equation 1, the killing parameter, *ω*, has a similar effect to the parameter λ. This is because it acts to delay the establishment of a plasmid-free population. The killing parameter in the bacteriocin model produces different dynamics to that for the TA model as it acts on all plasmid-free cells rather than just newly plasmid-free cells. A hierarchical Bayesian method for fitting the model, see Methods, allows these parameters to be determined from simulated and experimental plasmid loss curves, Supplementary Figure 2.

### Toxin-antitoxin and bacteriocin systems minimally impact growth rates

We produced a fluorescent reporter plasmid, constitutively expressing dasher GFP from the strong, constitutive 0XB20 promoter in order to place a burden on plasmid-bearing host cells and enable easy detection of cells still carrying the plasmid. From this plasmid we produced three further plasmids by inserting the plasmid-stability mechanisms: hok/sok, axe/txe and microcin-V. The plasmid construction is shown in Supplementary Figure 3. Using a microplate reader to collect growth data and a non-parametric Gaussian process method to fit the growth curves (Swain et al. 2016) we were able to quantify the effect of carrying the plasmid and the different plasmid-stability mechanisms. The growth rates were assayed in antibiotic free LB media to prevent any confounding effects on growth by the presence of kanamycin. The growth curves and model fits from which the maximum growth rates are estimated are shown in the Supplementary Figure 4. The maximum growth rates in Figure 2A show a significant load on the growth rate between EcN-Lux and the plasmid containing systems. The growth rate ratios, γ, calculated by dividing the plasmid-bearing growth rate distribution by that of the plasmid-free, Figure 2B, show growth rates for the plasmid-bearing cells of between 60 and 70% of the plasmid-free cells. There is some indication that axe/txe (EcN-Lux:pUC-GFP-AT) does have an increased burden. These results demonstrate the problem for plasmid stability; once the plasmid is dropped, the significantly higher growth in the plasmid-free strain cause the plasmid-bearing population to be outcompeted.

**Figure 2:**
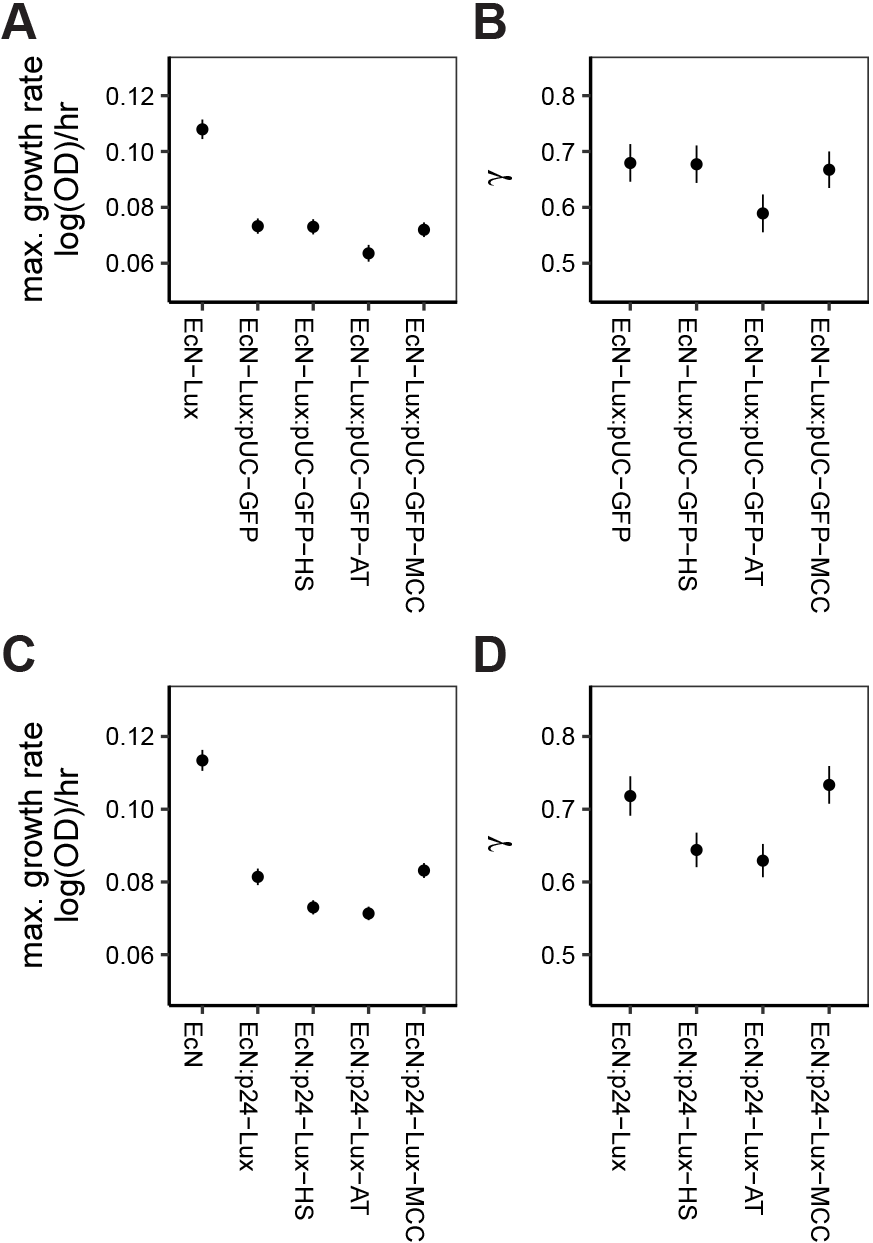
Growth rates and growth rate ratios for fluorescent and luminescent plasmids. (A) Maximal growth rates for the fluorescence plasmid in EcN-Lux were calculated by fitting a Gaussian process model to plate-reader data. (B) The ratio of plasmid-bearing growth rate to plasmid-free growth rate, calculated from (A), is equivalent to the plasmid-free doubling time over plasmid-bearing doubling time used in the plasmid-loss model. C) The Maximal growth rates and D) growth rate ratios for the luminescent plasmids in EcN. Points indicate the mean value and lines show the standard deviation of the mean.

We also produced luminescent reporter plasmids, Supplementary Figure 5, that constitutively express the *luxCDABE* operon from the phelp promoter (Riedel et al. 2007). The growth rates for the luminescence plasmid-bearing cells had to be assayed in LB media containing kanamycin to ensure plasmid maintenance. This was due to the instability of the control plasmid, p24-Lux. The growth rate results, Figure 2C and Supplementary Figure 6, show that there is a similar impact on growth rates compared to the fluorescence based plasmids.

### Both axe/txe and microcin-V outperform hok/sok in liquid culture

Flow cytometry data, Supplementary Figures 7-10, and an automated analysis pipeline (see Methods) were used to derive plasmid loss curves for the pUC-GFP based fluorescent plasmids in EcN-Lux over 37 daily passages (Figure 3A). For the control plasmid, without a PSK system (EcN-Lux:pUC-GFP), the plasmid-free cells start to become apparent after only ~ 4 passages, with the populations becoming entirely plasmid-free after ~ 12 passages. Figure 3A and Supplementary Figure 11 show that the toxin-antitoxin systems improve the stability of the plasmid, though the commonly used hok/sok (EcN-Lux:pUC-GFP-HS) only provides stability for ~ 16 passages, with one replicate beginning to drop as soon as passage 10 and one lasting until passage 22. The hok/sok system has a very high killing efficacy, failing to kill fewer than 5 × 10^−5^ new plasmid-free cells, as can be seen from the fitted model parameters in Supplementary Figure 12, but still loses out to competition from the few plasmid-free cells that escape the toxin. Strikingly, axe/txe (EcN-Lux:pUC-GFP-AT) remains stable throughout the experiment, clearly demonstrating its efficacy over the other systems. Because no plasmid loss is observed, the model fitting procedure cannot be applied as there is no information on the probability of plasmid loss, λ Finally, the results demonstrate that the microcin-V system (EcN-Lux:pUC-GFP-MCC) clearly outperforms the hok/sok system, although the results are varied, with four of the nine replicates remaining entirely plasmid-bearing for the length of the experiment and in five of the replicates we see a plasmid-free population taking over. The model determined efficacy of the bacteriocin is far below that of hok/sok, Supplementary Figure 12, though still manages to produce stable populations for a longer period.

**Figure 3:**
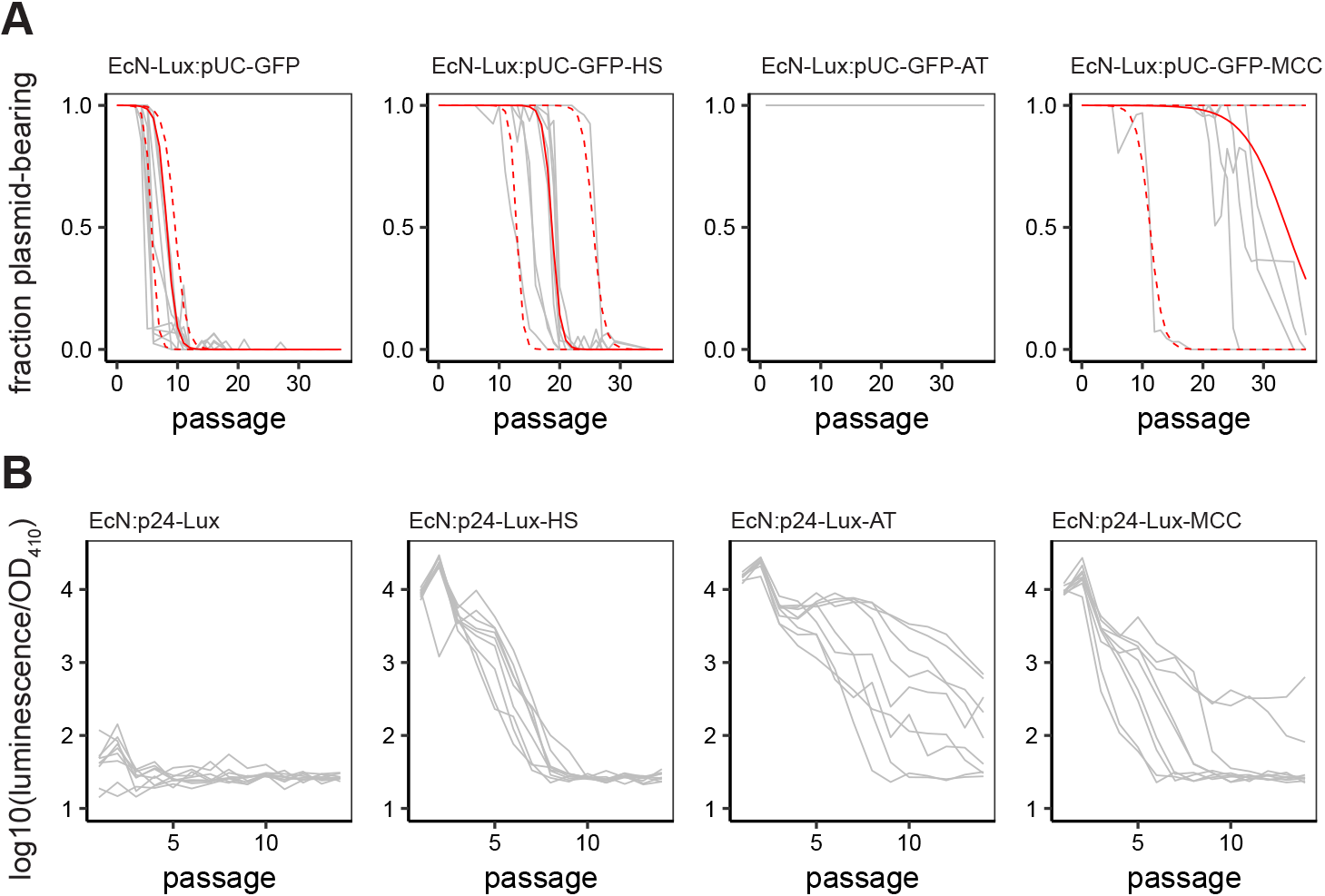
A) Plasmid loss curves for fluorescence based plasmids in EcN-Lux. The grey lines show the trajectories of 9 replicates for each strain. The solid red line shows the average model fit and the dashed red lines show the 95% confidence intervals of the model with posteriors from all replicates. B) Luminescence loss for the luminescence based plasmids in EcN. The grey lines show the trajectories of 9 replicates for each strain.

The raw flow cytometry data, Supplementary Figures 7-10, shows that when plasmid maintenance is enforced through the use of antibiotics the fluorescence levels remain stable across the full 37 passages for all but the microcin-V bearing strain. For the control plasmid and two TA system plasmids, this suggests that the plasmid copy number is stable and the fluorescent cassette on the plasmid has not mutated. It also indicates that samples being categorised as plasmid-free cells are not the result of loss-of-function mutations leading to a loss of fluorescence. This conclusion is further supported by the population level fluorescence data recorded using a microplate reader, Supplementary Figure 11. In contrast, we do see a reduction in fluorescence in the flow cytometry and microplate reader data for the microcin-V system. The flow cytometry measured fluorescence of the populations are homogeneous indicating that, although fluorescence levels are being reduced, it is unlikely that cells being categorised as plasmid-free have undergone a loss-of-function mutation to remove any fluorescence.

Plasmid-stability for the luminescent p24-Lux plasmid was then determined using the same growth protocol as for the fluorescent pUC-GFP based strains. However, measurements could only be carried out using population luminescence in a microplate reader rather than single cell fluorescence in a flow cytometer. Figure 3B shows that the control population (EcN:p24-Lux) has almost completely stopped luminescing by the end of the first passage. The two TA strains and the microcin-V carrying strain perform better than the control and rank in the same order as for the fluorescent plasmids. However, the luminescence loss occurs faster and even affects the axe/txe bearing strain. Even the samples in which the media contained kanamycin, enforcing plasmid maintenance, showed reductions in luminescence (Supplementary Figure 13), and in one of the EcN:p24-Lux replicates luminescence was completely lost.

### Microcin-V can restore a plasmid-bearing population

As we have seen with hok/sok, once plasmid-free cells arise in the population the toxin-antitoxin system has no way to save the plasmid-bearing population. Bacteriocins, however, are able to police the entire population due to the toxin being secreted into the environment. This means that when plasmid-free cells arise or if the culture becomes contaminated, the bacteriocin system can push the population back towards entirely plasmid-bearing.

Figure 4 shows that, when a population of TA plasmid-bearing cells is diluted with plasmid-free EcN-Lux, as stated above the plasmid-free population outgrows the plasmid-bearing population, further diluting it. However, when we dilute the EcN-Lux:pUC-GFP-MCC cells, they quickly kill the plasmid-free population and restore the population. In fact within 24 hours there are no plasmid-free cells detectable in all but one anomalous sample.

**Figure 4:**
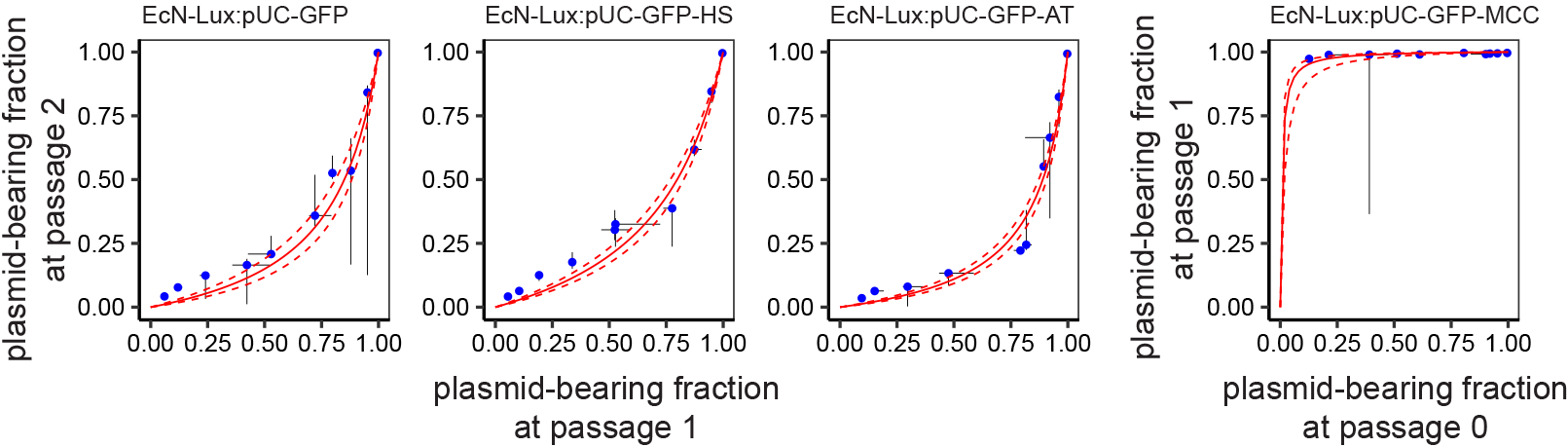
Plasmid-bearing strains are diluted with plasmid-free EcN-Lux at different ratios. The dilutions are sampled after 24 hours and show that the faster growth rate of the plasmid-free cells leads to further dilution of the plasmid-bearing population. In the case of EcN-Lux:pUC-GFP-MCC, however, the secreted bacteriocin allows the plasmid-bearing population to out-compete the plasmid-free cells. Blue points show the median of 3 replicates at each initial dilution; the bars show the minimum and maximum plasmid-bearing fraction at each passage. The solid red line shows the model fit and the dashed red lines show the 95% confidence intervals. Each replicate was passaged once initially to allow for the cells to acclimatise to the growth conditions. However, for the EcN-Lux:pUC-GFP-MCC replicates, most plasmid-free cells were dead after 1 passage so the data from passage 0 to passage 1 were used.

### Axe/txe and hok/sok successfully stabilises luminescent reporters *in vivo*

EcN strains containing either the p24-Lux-HS, p24-Lux-AT, p24-Lux-MCC or p24-Lux control constructs were intravenously injected into a mouse tumour model for *in vivo* characterisation (Danino et al. 2015). Once administered, the EcN strain was shown to colonise the tumours in the mouse flanks within 3 days (Figure 5A). The tumours were then excised 7 days after injection and a colony count was performed from the homogenised tissue to independently measure the stabilising effect of the axe/txe, hok/sok and microcin-V systems on the reporter construct. Figure 5B shows that without any stabilising system ~ 50% of the population had dropped the p24-Lux plasmid after intravenous injection and tumour colonisation. In comparison, the p24-Lux-AT was significantly stabilised (P = 0.0013) with nearly 100% of the population maintaining the plasmids after colonisation. The strain p24-Lux-HS also showed significant stability over p24-Lux control (P = 0.034) with ~ 80% of the population maintaining the plasmids after colonisation. The bacteriocin plasmid p24-Lux-MCC did not show a significant improvement over the control though the mean plasmid-bearing population was ~ 70. There was also no significant difference between the plasmid fraction bearing population in p24-Lux-AT and p24-Lux-HS.

**Figure 5:**
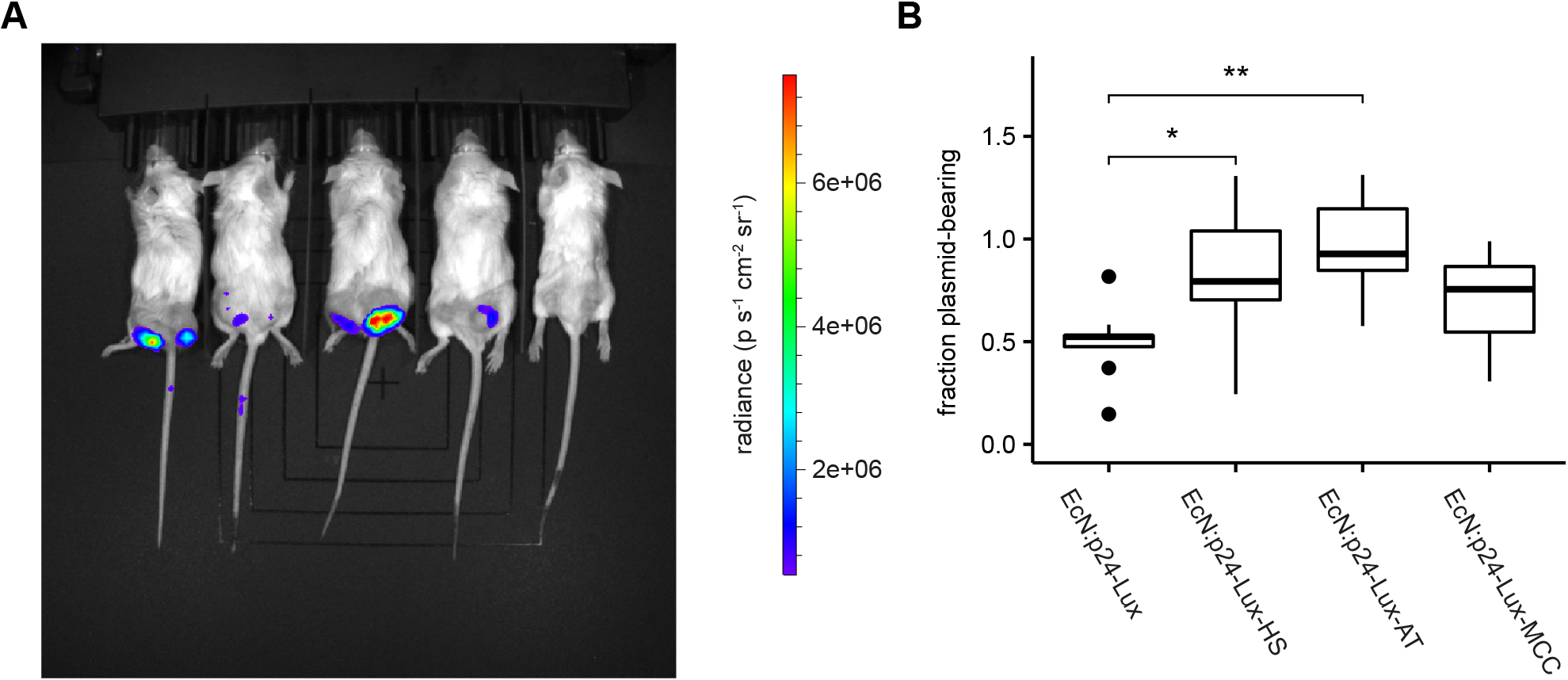
*in vivo* stability of plasmid-bearing EcN. (A) Representative image of plasmid produced luminescence after bacterial colonisation of implanted hind flank tumours. (B) Fraction of bacterial population remaining plasmid-bearing 7 days after colonisation, calculated from colony counts on selective and non-selective media performed in triplicate for each tumour. (*P=0.034, **P=0.0013 in the Mann-Whitney U test for p24-Lux n=9 tumours, p24-Lux-HS n=7 tumours, p24-Lux-AT n=8 tumours, p24-Lux-MCC n=5 tumours)

## Discussion

We developed an automated flow cytometry pipeline, mathematical model, and Bayesian inference procedure to quantify the efficacy of PSK systems stabilising fluorescent and luminescent reporter constructs *in vitro* and *in vivo*. We used this pipeline to demonstrate that two new PSK systems in *E. coli* Nissle, axe/txe and microcin-V, can outperform the more commonly used hok/sok system. Our method for Bayesian parameter estimation is able to accurately infer parameters from simulated data, and, where plasmid loss occurs, can be used to estimate growth rate differences, plasmid loss rates, and post-segregational killing efficacy.

As expected, plasmids expressing fluorescent and luminescent proteins were shown to reduce the growth rate of EcN significantly, but there was minimal extra burden produced from the toxin-antitoxin and bacteriocin systems. Our plasmid stability assays with both the fluorescent and luminescent plasmids showed that axe/txe provides greater plasmid stability than hok/sok. This is an interesting result as hok/sok is an *E. coli* native TA system whereas axe/txe was found in the gram-positive *E. faecium*. Further, hok/sok is bactericidal (Faridani et al. 2006) whereas axe/txe is only bacteriostatic (Grady & Hayes 2003). One might expect that the plasmid-free persister cells created from the txe toxin would begin to divide at a later passage and cause the plasmid-bearing population to become diluted but this did not appear to happen during the 37 passages of the fluorescent strains.

The results for microcin-V show far more variability in plasmid-stability than for the TA systems. This is likely to be due to the fact that bacteriocins have not evolved as plasmid stability mechanisms but rather as competition mechanisms. The dynamics of gene regulation within the microcin-V cassette have not been determined other than to show regulation through iron depletion (Boyer & Tai 1998). Supplementary Figure 14 shows potential transcription factors involved in microcin-V regulation but further empirical examination is needed to determine the conditions under which microcin-V is expressed. It may be necessary to replace the native promoters with a better understood regulatory system in order to achieve the type of population policing that was envisioned in Figure 1E. Further, we have shown that the of carrying the microcin-V system leads to reduced protein expression from the plasmid after several generations. This may hinder its use for prolonged applications.

Although microcin-V does not perform as well as axe/txe over the length of the plasmid-stability experiments, we have shown that over a single passage of 24 hours the bacteriocin is able to push a diluted population back to entirely plasmid-bearing. For this reason, microcins may be particularly effective in industrial fermentation applications where the environment is relatively stable compared to the *in vivo* conditions encountered in clinical use.

A major focus of synthetic biology over the coming years will be to push proof-of-principle systems into real-world applications. A big limitation for industrial and clinical applications adopting synthetic biology tools is the use of antibiotic selection and moving away from these laboratory-based systems will be critical. Our developed experimental and modelling approach can be used to characterise new systems for industrial applications and novel therapeutics. By inferring information about the relative growth rates and killing efficiency, a more quantitative viewpoint of plasmid stability mechanisms can be achieved, enabling the targeting of PSK systems to specific applications.

## Experimental Procedures

### Author Contributions

A.J.H.F., T.O, A.D., L.R, and O.V., designed and performed experiments. A.J.H.F., T.O, T.D. and C. B conceived this study, analysed the data, discussed results, and wrote the manuscript.

## Acknowledgements

We thank Professor Kenn Gerdes for sending the pKG1022 plasmid (now at the University of Copenhagen), Dr Finbarr Hayes at The University of Manchester for sending the pREG531 plasmid, Professor Roberto Kolter at Harvard for the pHK11 plasmid and Dr Esteban Martinez-Garcia at CNB-CSIC for the pSEVA246 plasmid. A.J.H.F is funded through the UCL CoMPLEX doctoral training centre. T.O. is funded through the BBSRC LIDo doctoral training partnership. L.R. is funded jointly through a BBSRC studentship and a Microsoft Research Scholarship. T. D. was supported by an NIH R00 (CA197649) and DoD Era of Hope Award (BC160541). C.P.B. is supported through a Wellcome Trust Research Career Development Fellowship (097319/Z/11/Z). We thank C. Coker and J. Zhang in the Danino laboratory for assistance with mouse experiments.

